# An exploration of the SARS-CoV-2 spike receptor binding domain (RBD) – a complex palette of evolutionary and structural features

**DOI:** 10.1101/2020.05.31.126615

**Authors:** Dwipanjan Sanyal, Sourav Chowdhury, Vladimir N. Uversky, Krishnananda Chattopadhyay

## Abstract

SARS-CoV-2 spike protein (S) is associated with the entry of virus inside the host cell by recruiting its loop dominant receptor binding domain (RBD) and interacting with the host ACE2 receptor. Our study deploying a two-tier approach encompassing evolutionary and structural analysis provides a comprehensive picture of the RBD, which could be of potential use for better understanding the RBD and address its druggability issues. Resorting to an ensemble of sequence space exploratory tools including co-evolutionary analysis and deep mutational scans we provide a quantitative insight into the evolutionarily constrained subspace of the RBD sequence space. Guided by structure network analysis and Monte Carlo simulation we highlight regions inside the RBD, which are critical for providing structural integrity and conformational flexibility of the binding cleft. We further deployed fuzzy C-means clustering by plugging the evolutionary and structural features of discrete structure blocks of RBD to understand which structure blocks share maximum overlap based on their evolutionary and structural features. Deploying this multi-tier interlinked approach, which essentially distilled the evolutionary and structural features of RBD, we highlight discrete region, which could be a potential druggable pocket thereby destabilizing the structure and addressing evolutionary routes.

## Introduction

The 2019 novel coronavirus has spread worldwide, becoming a pandemic by affecting more than 200 countries and becoming one of the worst infections in the recent times. Efforts at multiple levels are currently ensuing to effectively combat the disease spread and to come up with some potential preventive or therapeutic solutions. Majority of the therapeutic strategies are aimed at blocking the virus entry into the host cells, which is primarily driven by the SARS-CoV-2 spike (S) protein and its interaction with the angiotensin-converting enzyme 2 (ACE2) that is located at the outer membranes of cells in the lungs, arteries, heart, kidney, and intestines and serves as a SARS-CoV-2 receptor. The coronavirus entry into the host cell is a complex, multi-step, highly orchestrated process, which involves multiple processing stages of the S protein. The entire spike protein S is cleaved into S1 and S2 subunits, with S1 further participating in receptor recognition by recruiting its receptor binding domain (RBD). The S protein, being a key target for vaccines, therapeutic antibodies, and diagnostics, has been studied by various groups using different techniques, and the loop dominant RBD region of the protein has been reported to be the key in the receptor binding (1). The residues in RBD that are not directly involved in interaction with receptor but impose structural constraint on the motif/region (receptor binding region) may serve as druggable pockets for small molecule binding, depending upon their evolutionary traits and impact on global structure of the protein. Hence, we truncated and removed this domain (RBD) from the overall protein structure (**Figure 1**) and critically analyzed various aspects of Spike-RBD, which stands immensely important for comprehensive understanding of the importance of this protein.

**Figure 1:**
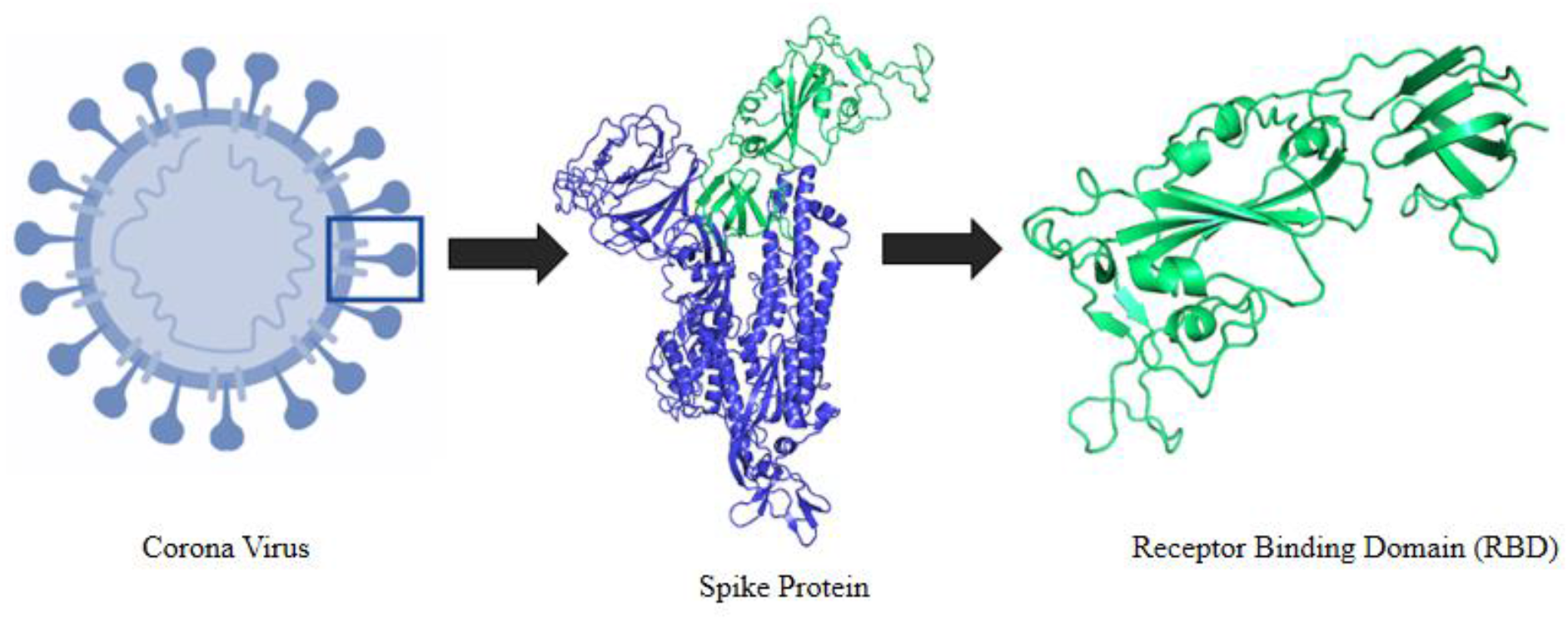
Schematic representation of the selection logistics of the protein and protein segment. It shows an animated figure of the 2019 novel coronavirus. Segment presented in green cartoon refers to the receptor binding domain (RBD) of the spike protein.

We have resorted to a two-tier approach in our study encompassing evolutionary and structural fabric of RBD, which is a 273 amino acid segment of the whole Spike protein stretching from residue 319 to 591. In our study, especially in the sequence analysis segment, we have referred this residue stretch from 319 to 591 as positions 1 to 273. RBD, after the initial processing, docks into the binding cleft of the ACE2 receptor of host cells and thereafter initializes the virus entry process into the host cell. Our evolutionary tier of the current study was primarily focused on the sequence space of the RBD. The sequence space recreation clearly portrayed the closest RBD relatives and helped understanding the evolutionary constraints. Understanding evolutionary constraints provides a glimpse into the evolvability scope of a protein and helps conceiving an anti-evolution therapeutic strategy, which could impact the evolutionary trajectory. We observed sections of RBD, which were evolutionarily conserved as quantified by Shannon’s entropy and the regions of co-evolutionary importance. Unlike sequence conservation, which gives a static snapshot of positions, which were critical and hence not subjected to variation, co-evolutionary analysis provides information on the positional interdependencies and thereby helps recreating a dynamic picture of residue-residue dependence, which, in turn, is a key to stabilize protein structure and explain the functionalities of a protein. We chose to use both sequence conservation and co-evolution signals from the RBD sequence space to better understand the constraints in the RBD sequence space and the RBD evolutionary dynamics. Using mutual information (MI) values which correspond to the positional interdependencies we observed region 121 to 180 (i.e., 439 to 498 w.r.t the S-protein) of the RBD having maximal co-evolutionary signal. We further used this sub-space (121 to 180) for deep mutational scan using epistasis model to generate a mutational landscape, which could potentially present a picture of the substitution mutation tolerance at each of the sites comprising the sub-space. We searched for all possible naturally occurring mutations, which were observed at the sites, which have been reported to interact with the host ACE2 receptor by using the sequence information derived from the closest relatives as obtained from the phylogenetic analysis. Further, we computed their impact using statistical energy computed from the deep mutational scan. Finally we correlated the residues which were observed to be critical in co-evolution with their mutational tolerance.

We used the RBD structure (319 to 591 w.r.t S-protein) for structure network analysis and Monte Carlo simulation studies to respectively dissect the whole protein into sub-structures (referred as sub-blocks or SB in the article) based on their pair-wise interaction and local dynamics as quantified using root-mean-square fluctuations. Our analysis revealed how residues are clustered based on their pair-wise interactions and the region of the protein with the highest local fluctuation. Our docking data further provided an insight into these highly fluctuating local regions, which were in the close vicinity to the residues L455, F486, Q493, S494, and N501 making up the RBD binding cleft. Unlike the previous reports which commented upon the importance of the loops in the RBD in general, we specifically pin-pointed this stretch and hypothesized that it is extremely critical in conferring local conformational flexibility leading to an effective interaction with ACE2 receptor.

Finally, to have a comprehensive insight into the RBD from both evolutionary and structural perspective, we plugged information from both tiers of our study and applied fuzzy logic principles to see, which regions of the protein have maximal overlap in terms of their evolutionary and structural features. For this we selected only those structure blocks (SBs), which had maximum quantified evolutionary (conservation and co-evolution) and structural (based on properties viz. hydropathy, higher order structure etc.) information. This cumulative approach helped holistically understand region of RBD, which if destabilized (which we believe to be a potential therapeutic strategy), would thoroughly impact the evolutionary routes and biophysical integrity (both structural and receptor-recognition centric function) of the protein. Thus resorting to an inter-linked multi-tier approach, we thereby project segments of RBD, which could be a potential therapeutic target to not only impact the structure but also disrupting the RBD evolutionary trajectory and hence a potential anti-evolution strategy.

## Results

### Sequence space analysis

Our sequence space analysis revealed that Spike RBD of SARS-CoV-2 has high degrees of sequence identity with other members of the spike glycoprotein family of Bat coronavirus origin. The phylogenetic tree as constructed using the neighbor-joining method provides a glimpse of the 15 closest proteins retrieved from the sequence space of Spike RBD (**Figure 2**). This analysis provided an insight into the sequence divergence and closest relative of the Spike RBD, which, as revealed by this phylogenetic analysis, happens to be Spike protein (Fragment) n=1 Tax=Bat coronavirus TaxID=1508220 (UniRef90: A0A5H2WUD2).

**Figure 2.**
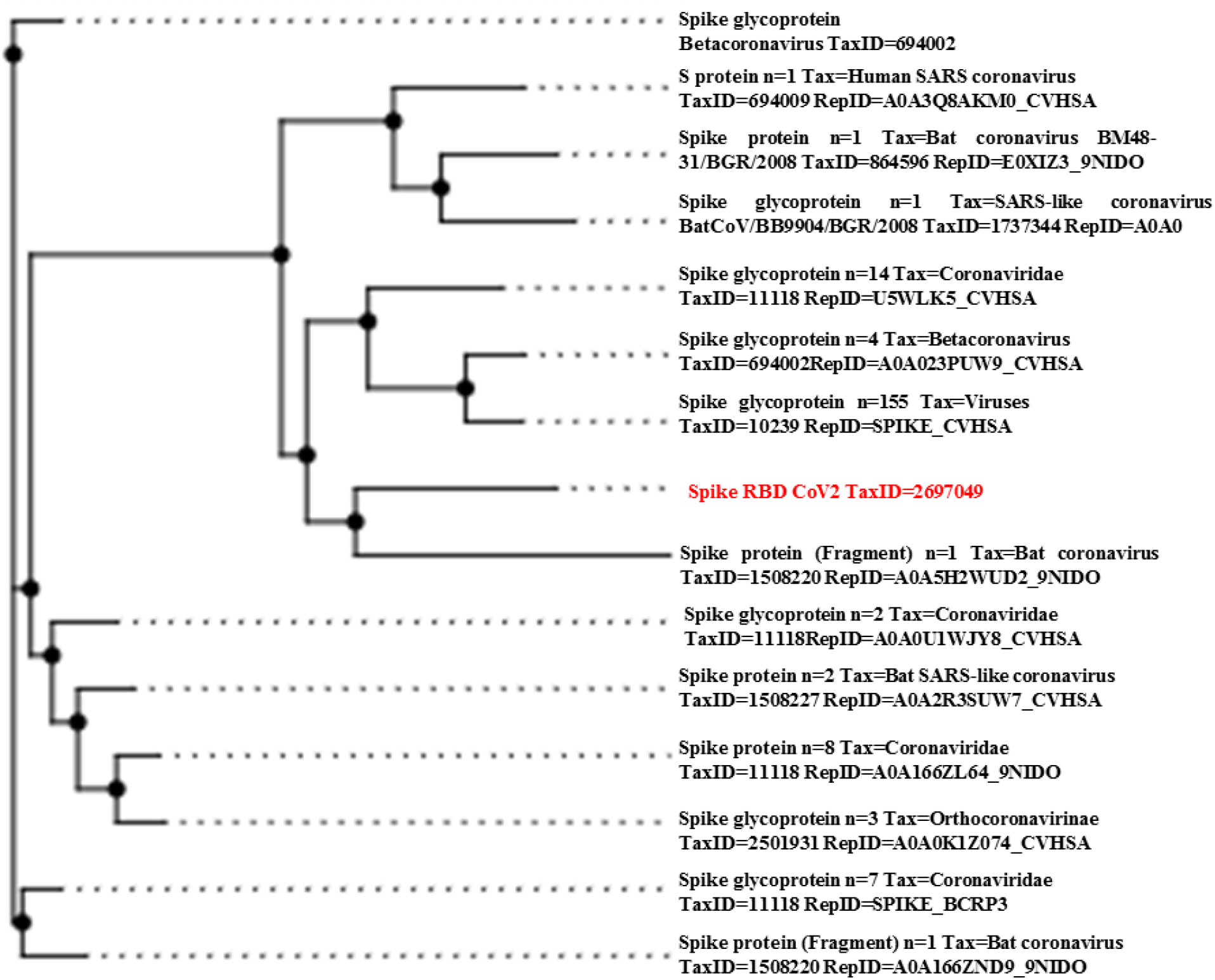
Phylogenetic tree of the Spike RBDs showing 15 closest relative of the RBD from SARS-CoV-2 as retrieved from the sequence alignment results.

Next, we analyzed the preservation of the intrinsic disorder predisposition within the amino acid sequences of the 15 evolutionary closest Spike RBDs. Results of this analysis are summarized in **Figure 3** showing rather close similarity of the overall shapes of the disorder profiles of these domains. One can clear observe comparable patterns, where more flexible or even intrinsically disordered regions (i.e., regions with the disorder scores between 0.2 and 0.5 and above 0.5, respectively) are interspersed among more ordered segments. Many of these flexible/disordered regions correspond to the RBD loops. **Figure 3** also demonstrated that the largest variability in the per-residue disorder predisposition is observed for the C-tail of RBDs and their centrally located, 60-residue-long region (residues 121-180).

**Figure 3.**
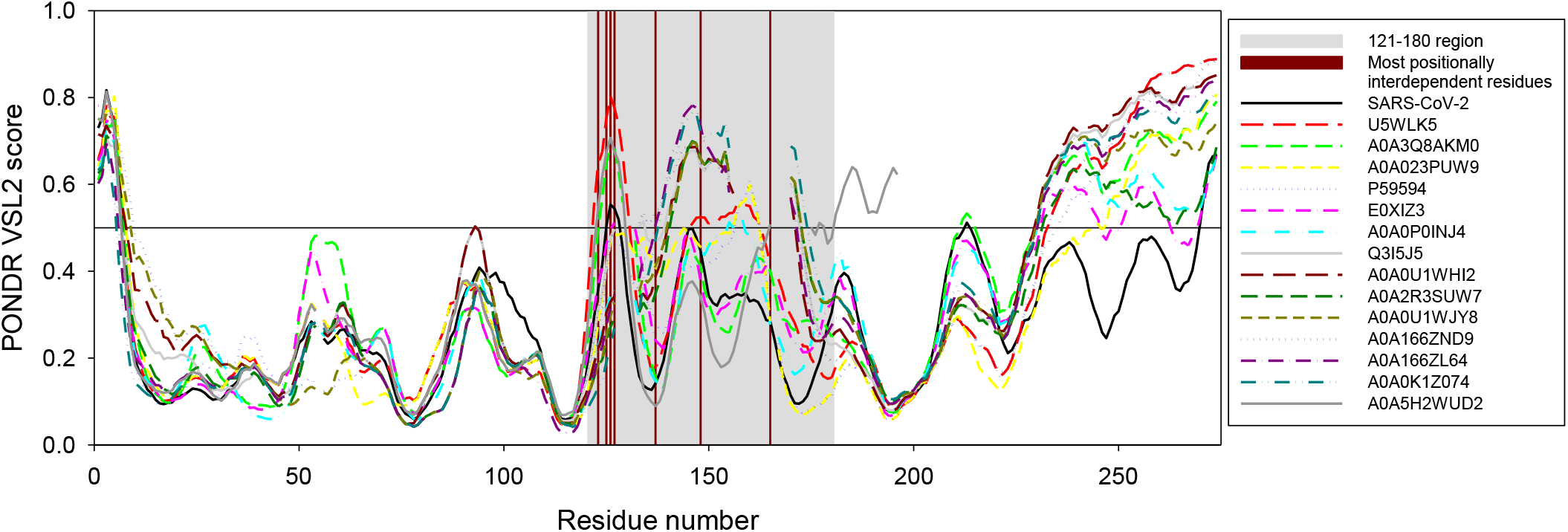
Per-residue intrinsic disorder predisposition of the 15 most evolutionary similar Spike RBDs evaluated by PONDR^®^ VSL2(2). Breaks in the disorder profiles correspond to the breaks in sequence alignments. In this analysis, disorder scores between 0.2 and 0.5 indicate flexible regions, whereas regions with the disorder scores ≥0.5 are expected to be intrinsically disordered. Position of the 121-180 region corresponding to the area with the highest co-evolutionary signal and thus with the maximal positionally interdependent residues is shown by light gray shading. Dark red bars shows position of residues which are most positionally interdependent (123, 125, 126, 127, 137, 148, and 165).

**Figure 4A** provides a sequence conservation profile of Spike RBD. The conservation score at a site corresponds to the site’s evolutionary rate. The rate of evolution is not constant among amino the sites. Some positions evolve slowly and are commonly referred to as “conserved”, while others evolve rapidly and are referred to as “variable”ate variations correspond to different levels of selection acting on these sites. For example, in proteins, the purifying selection can be the result of geometrical constraints on the folding of the protein into its 3D structure, constraints at amino acid sites involved in enzymatic activity or in ligand binding or, alternatively, at amino acid sites that take part in protein-protein interactions. From the Shannon’s entropy contour as a function of residue index (**Figure 4A**), it could clearly be suggested that the evolutionarily conserved residues having zero shannon entropy measurement were scattered throughout the RBD sequence space. With increase in the Shannon’s entropy values, extent of conservation decreased with the increment in propensity to be more interdependent. In a broad patch of region from around 122 to 232 (i.e., residues 440 to 550 with respect to the S-protein) entropy profile showed some spike (high values) indicating the lesser extent of conservation (**Figure 4A**).

**Figure 4.**
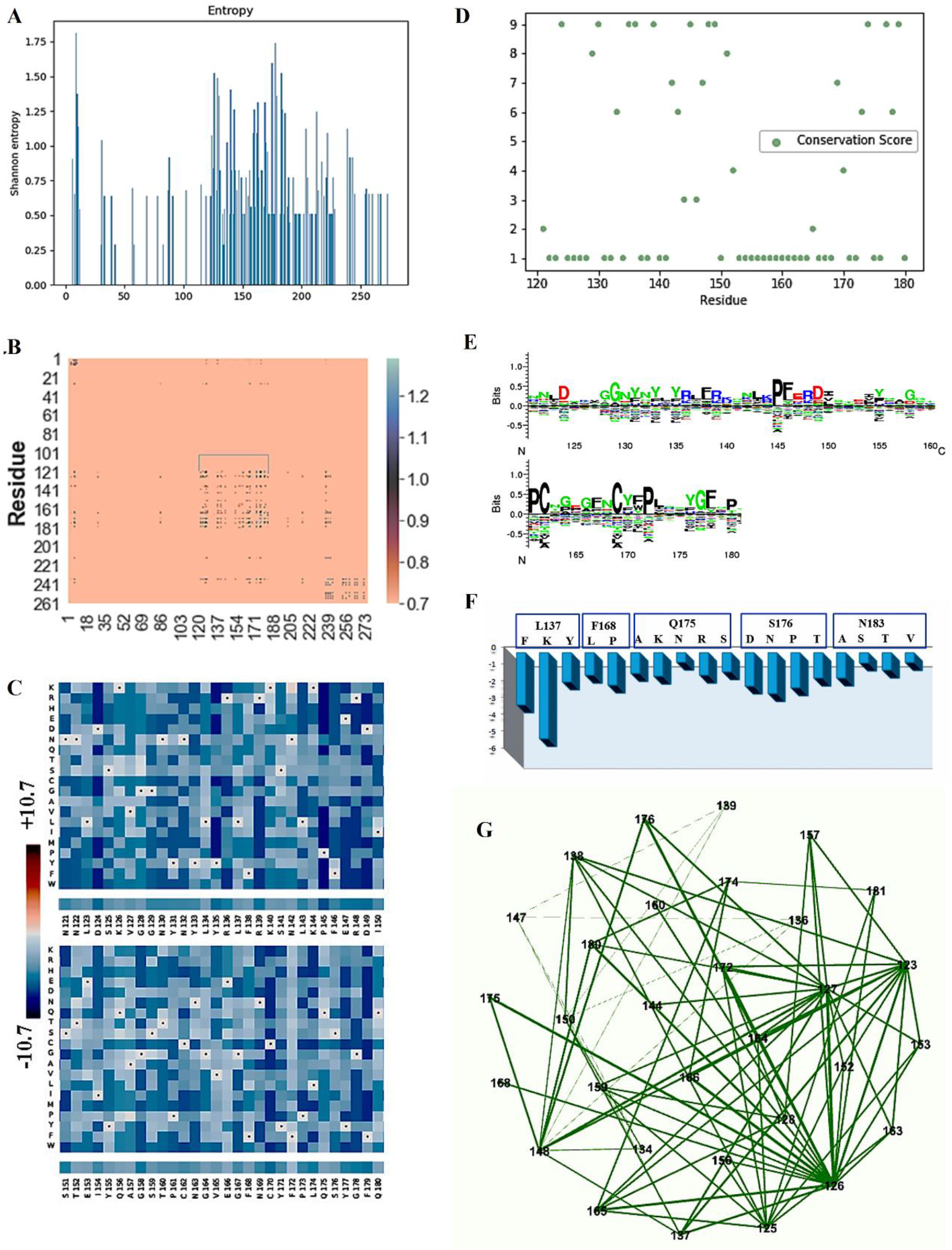
Sequence space exploration of the spike RBD reveals critical areas, which are under the evolutionary constraints. (**A**) Shannon Entropy profile of the RBD sequence space. (**B**) Heat-map showing co-evolution as quantified using mutual information values, in the RBD sequence space. Colour bar in the left shows the gradient of MI values in the co-evolving sites. Bracketed (black) segment shows the patch with highest co-evolution. (**C**) Mutational landscape as generated from the deep mutation scan is shown as a heat-map with color bar in the left showing the gradient of statistical energy. The bottom bars below each of the sub maps depict the sequence information for the spike RBD. (**D**) Conservation profile of the spike RDB as calculated by Bayesian method. (**E**) Sequence logo depicting the RBD sequence space as calculated using Kullback-Leibler method. (**F**) Bar plots depicting differences in statistical energy (dE) profiles upon substitution mutations with residues derived from variations observed among the closest relatives at positions known to be critical in RBD-ACE2 interaction. (**G**) Fruchterman-Reingold graph generated using MI values showing the network of co-evolving sites in the RBD sequence space.

It is to be noted that sequence space constraints are primarily dictated by positional conservation and positional inter-dependencies; i.e., co-evolution. Sequence conservation provides a static snapshot capturing position-conservation and vastly under-represents the evolutionary dynamics as position-position interdependencies, which in turn regulate inter-residue interactions in the 3-D space of a protein structure, remain completely uncharted. After gaining an insight into the sequence conservation profile, we went ahead to understand the sequence co-evolution profile resorting to mutual information (MI) calculations on the aligned sequences. A high MI value corresponds to the higher positional inter-dependency and hence higher co-evolutionary signal from the sequence space. To capture highest co-evolution signals, which in turn could reliably dictate the sequence space constraint, we used a threshold of MI values > 0.7 and filtered those signals to generate the co-evolutionary matrix (**Figure 4B**). The dotted area in the residue-residue co-evolutionary matrix (also bracketed in the heat-map) reflects the sub-space in the sequence space of spike-RBD which has the highest co-evolutionary signal and hence maximal positionally inter-dependent residues. This region, which spans from position 121 to 180 (439 to 498 w.r.t. to whole protein) was further subjected to deep-mutation scans using Epistasis model to predict possible effects on the overall stability in the face of substitution mutation. Given the fact that viral genomes are highly prone for mutation events, which in turn shapes their evolutionary fitness, this current exercise utilizing deep mutation scan and generating mutational landscape was therefore extremely critical in terms of understanding the positional tolerance and its concomitant effect on biophysical fitness (both on structure and receptor-recognition centric function). Mutational landscape (**Figure 4C**) shows a relatively high position-wide residue tolerance, which in our calculation is a function of -∆E (statistical energy) with particularly higher tolerance at positions at 124, 135, 139, 145, 158, and 170 (442, 453, 457, 463, 476 and 488 respectively w.r.t the S-protein), where specific residue-based substitution is likely to confer higher stability. Intensity of blue colour in the boxes (**Figure 4C**) represents the degrees of stability being conferred upon substitution in the background of the current sequence, as guided by the colour bar on the left which scales the range of ∆E. This predictive mutational landscape based on epistasis model gives critical clues as to which substitutions could the sequence space potentially witness under the sequence space constraints.

We further deployed Bayesian method to dissect the position stretch spanning from 121 to 180 (i.e., from 439 to 498 in case of RBD), which represents maximal co-evolution signal and calculate the conservation score. Conservation score reflects (**Figure 4D**) that this stretch is characterized by an interesting interplay of conservation and variation. It is interesting to note that positions spanning from 150 to 164; i.e., 468 to 482 w.r.t the RBD (save 151 and 152) do show very low conservation, and, when compared to the mutation landscape, these regions also exhibit high tolerance. On the other hand, the positions with higher conservation (spanning from 143 to 149; i.e., 461 to 467 the S-protein) have relatively stricter bias towards to substitution mutations and lesser tolerance (**Figure 4C**). We thereby provide a unified picture of how sequence conservation could be potentially addressed using deep mutation scan approach and is insightful in the current context of spike-RBD sequence space.

We went on to generate the statistical energy profiles of the most commonly encountered substitution mutations in the sequence space for those residues, which are reported to participate in ACE2 receptor recognition. A sequence space logo as generated using Kullback-Leibler principle shows the residue variations and the pre-dominant residue occurrences in the sequence space (see **Figure 4E**). Residue substitution information was retrieved from those evolutionary trajectories, which were closest and were used for generating the phylogenetic tree (**Figure 2**). Statistical energy profile (**Figure 4F**) gives a quantitative idea as to which mutations would vastly stabilize the protein. Save the lysine substitution at position 137 (which designates residue 455 in the S-protein) others mostly show comparable extent of stability and hence explain why they could occur in the other close relatives of Spike-RBD yet retaining the same structural integrity.

We further utilized the co-evolutionary signal to generate a weighted matrix and went on to apply graph theoretical approach to map the highest co-evolving residues. Using the Fruchterman-Reingold graph layout (**Figure 4G**) we observe that positions 123, 125, 126, 127, 137, 148, and 165 (441, 443, 444, 445, 455, 466 and 483 respectively w.r.t. S-protein) represent the nodes with maximum edges; i.e., many residues are positionally interdependent. It is worth noting that these positions (except 148; i.e., 466 w.r.t. S-protein) also exhibited higher tolerance in the mutational landscape. As these positions are evolutionarily coupled with multiple residues, a mutational tolerance provides a dynamic evolutionary fabric which would tolerate mutations at other coupled positions and yet evolve in current constraints in the sequence space. **Figure 3** shows that many of these residues are located within intrinsically disordered regions.

Our comprehensive sequence space analysis utilizing information theory, deep mutation approach, and graph theoretical analysis captures specific positions, which are extremely critical, and shapes the constraints in the sequence space of spike-RBD. Our results have shown the interesting aspects of position spanning from 121 to 180 (439 to 498 w.r.t S-protein), which captures the co-evolutionary picture of spike-RBD and depicts the evolutionary dynamics of the spike RBD.

### Structural analysis

In order to unravel the internal arrangement and inter-dependency of the residues in terms of their pairwise interaction, structure network analysis was deployed (3). By generating an all-residue network coupled with community clustering, the evolutionarily coupled co-varying patches in the RBD were further analyzed. The residues of the RBD of the protein were found to be split into 30 sub-blocks (SBs); i.e., they were distributed through the space by forming 30 clusters (**Figure 5A**). Among these clusters, mainly 5 were found to be comprised of large number of residues. Most of the remaining clusters were represented by a single residue, in some cases 3 or 4 residues were housed together (SI). Sub-block 2, 6, 12, 17, and 21 were found to be comprised of large number of residues (**Figure 4**). Also, due to their dense connection with other SBs, these clusters were selected for study in detail to understand how the residues in those clusters were significant in terms of evolutionary features, receptor binding, as well as local and global motion.

**Figure 5.**
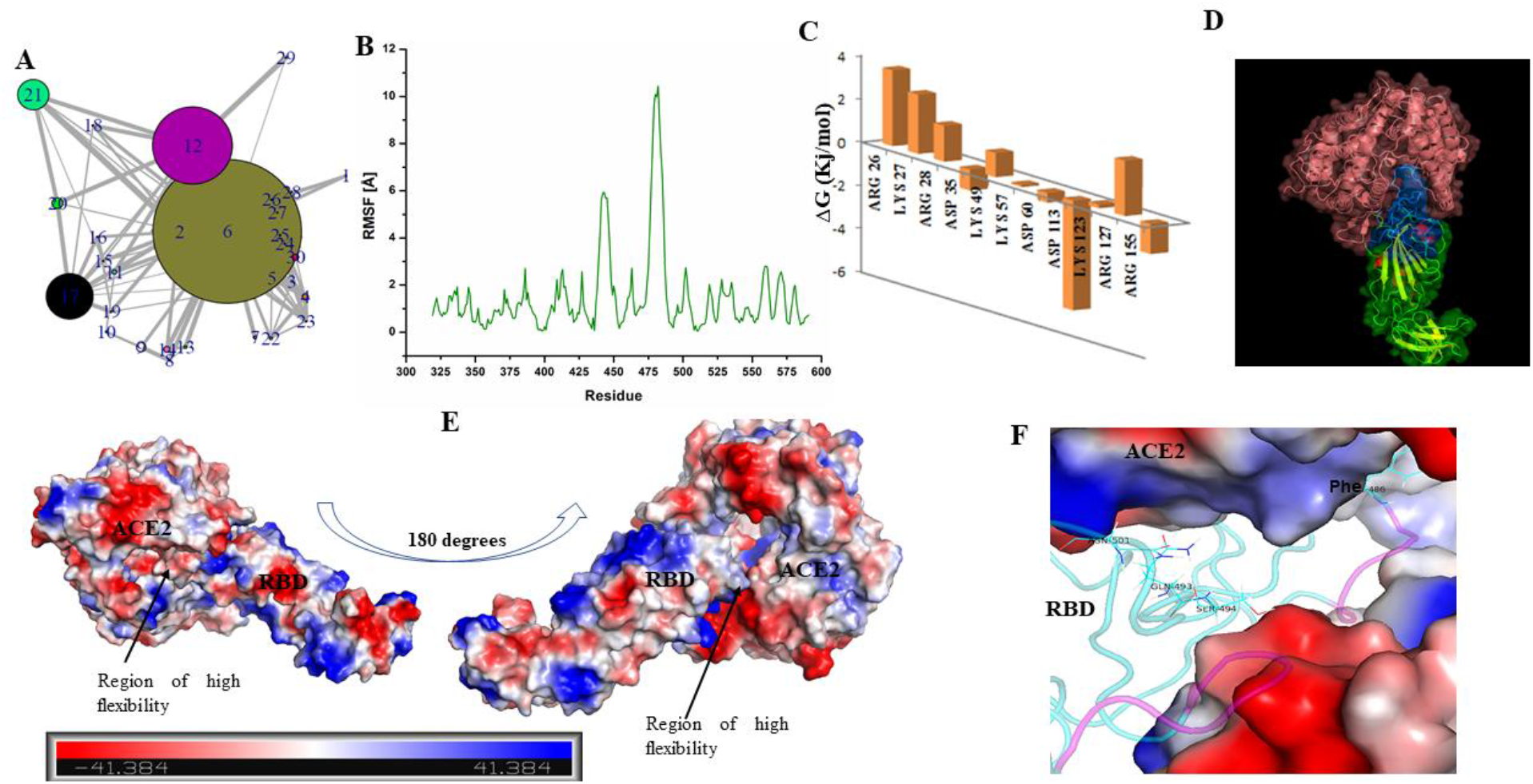
Structure of RBD has loop dominant region, which plays critical role in ACE2 receptor recognition and associated conformational dynamics. (**A**) Structure Network plot of spike RBD. (**B**) Root mean square fluctuations (in Angstrom) for the spike RBD as retrieved from Monte Carlo simulation. (**C**) Bar plots showing the effects of destabilizing mutations. (**D**) Docked complex of Spike RBD-ACE2 is shown in surface representation with cartoons representing the protein secondary structures. ACE2 receptor is shown in pink while RBD is shown as blue and green surfaces. Blue surface refers to the region in the RBD which participates in ACE2 receptor binding. (**E**) Docked complex is shown with surface charge representations. Region of high flexibility as observed from MC simulation is marked in the docked complex. Scale bar in the bottom refers to the gradient of surface charge as observed in the docked complex. (**F**) RBD-ACE2 interaction pocket (ACE2 is shown as surface and RBD in cartoon) is shown with purple coloured loops indicating substructure with high flexibility (as predicted from RMSF).

According to the docking output, receptor was observed to have a cleft, which accommodated the interacting loop region of the RBD (**Figure 5D**). Residue patches from 28/346 to 34/352, 97/415 to 102/420 and the loop segment from residue 126/444 to residue 182/500 were witnessed to be extremely decisive for the surface overlap. Monte Carlo simulation was performed with the Spike RBD to understand the dynamics associated with the Spike receptor binding domain. RMSF profile as retrieved from our simulation studies showed residue stretch 159/477 to 166/484 and 123/441 to 127/445 having a very high fluctuation level of >6 Å (**Figure 5B**). It is interesting to note that this region of high RMSF, which indicates high local dynamics, corresponds to two loop regions (highlighted in magenta, see **Figure 5F**) in the close vicinity to the residues associated with the ACE2 receptor recognition. This region is also predicted to be highly disordered (see **Figure 3**), and we believe that this can confer allostery and conformational flexibility guiding effective receptor binding. As the bound complex formation is not entirely governed by opposite surface charges (**Figure 5E**), we hypothesize that local flexibility at stretches 123/441 to 127/445 and 159/477 to 166/484 is extremely critical in conferring conformational flexibility associated with a rather efficient receptor binding.

It is further interesting to note that the segment of Spike-RBD, which resides in the close proximity to the ACE receptor complex (highlighted in blue, **Figure 5D**) strikingly overlaps with the position stretch of high co-evolution signal as retrieved from our evolutionary studies. A highly clustered abundance of mutually interdependent co-evolving residues clearly portrays the deep evolutionary importance of this region. The stretch has a very low structural organization and is pre-dominantly loop-dominant, which presumably provides extreme structural malleability in receptor recognition and associated allostery, and, hence its biophysical fitness. Loop regions in a protein further provide the ability to adopt transient context dependent conformations (4–6). Previous reports have highlighted the importance of residue L137/455, F168/486, Q175/493, S176/494 and N183/501 in receptor recognition (7). Our simulation-guided approach further establishes two additional sub-stretches, which potentially regulate effective interaction between the spike protein and its ACE2 receptor.

The TKSA-MC method refers to the residues which has a destabilizing contribution to the protein native state(8). The algorithm calculates the protein electrostatic energy, taking into account the contribution of each residue with polar charged side chain. The bars in the bar-plot show the charge-charge energy contribution of residues, which are ionisable with respect to the protein native state stability. The bars on the positive y-axis refer to the residues to be mutated to enhance the protein thermal stability. As per Ibarra-Molero model, residues with unfavorable energy values show ∆G_qq_ ≥ 0 and are exposed to solvent with SASA ≥ 50%(9). In the spike RBD protein, residues, which got selected in the destabilization simulation study, were predominantly outside the evolutionarily constrained region of 121 to 180. Residues 123 and 127 (designating position 441 and 445 in the S protein) occurring inside the evolutionarily constrained stretch have interestingly a high tolerance towards the substitution mutations in the deep mutation landscape with a probable indication of better fitness and cross-validated by TKSA-MC analysis.

### Comparison between Sequence and Structure index

After capturing both sequence space and structure-based aspects of RBD, we went on to identify a stretch in the structure, which contains the highest evolutionary and structural information and could be a key for regulation of the evolutionary and structural trajectory of the protein. As discussed in the methods section, we specifically focused on selected evolutionary and structure parameters, which could reliably project evolutionary and structural information. We dissected the whole RBD into discrete structure blocks based on structure-network analysis and plugged evolutionary and structural information on to these blocks.

The cluster dendrogram (**Figure 6A**) quantitatively represents the evolutionary and structural information associated with the top structure-blocks (the ones with significant evolutionary and structural information) and their extent of similarity. From the tree analysis (**Figure 6A**), it was observed that SB21 shared similarity with SB12 in terms of the defined evolutionary and structural index. SB17 demonstrated similar characteristics with the previous two clusters with higher extent of overlap. Similarly, SB6 was found to resemble in properties SB2 (**Figure 6A**). Here, the imposed evolutionary property indexes displayed the propensity of the protein segments (SBs) towards conservation and interdependency. Defined structural properties dictated the contribution of the residues and, in turn, the clusters. The dendrogram was a representation of integral correlation between the SBs in terms of the above-mentioned properties.

**Figure 6:**
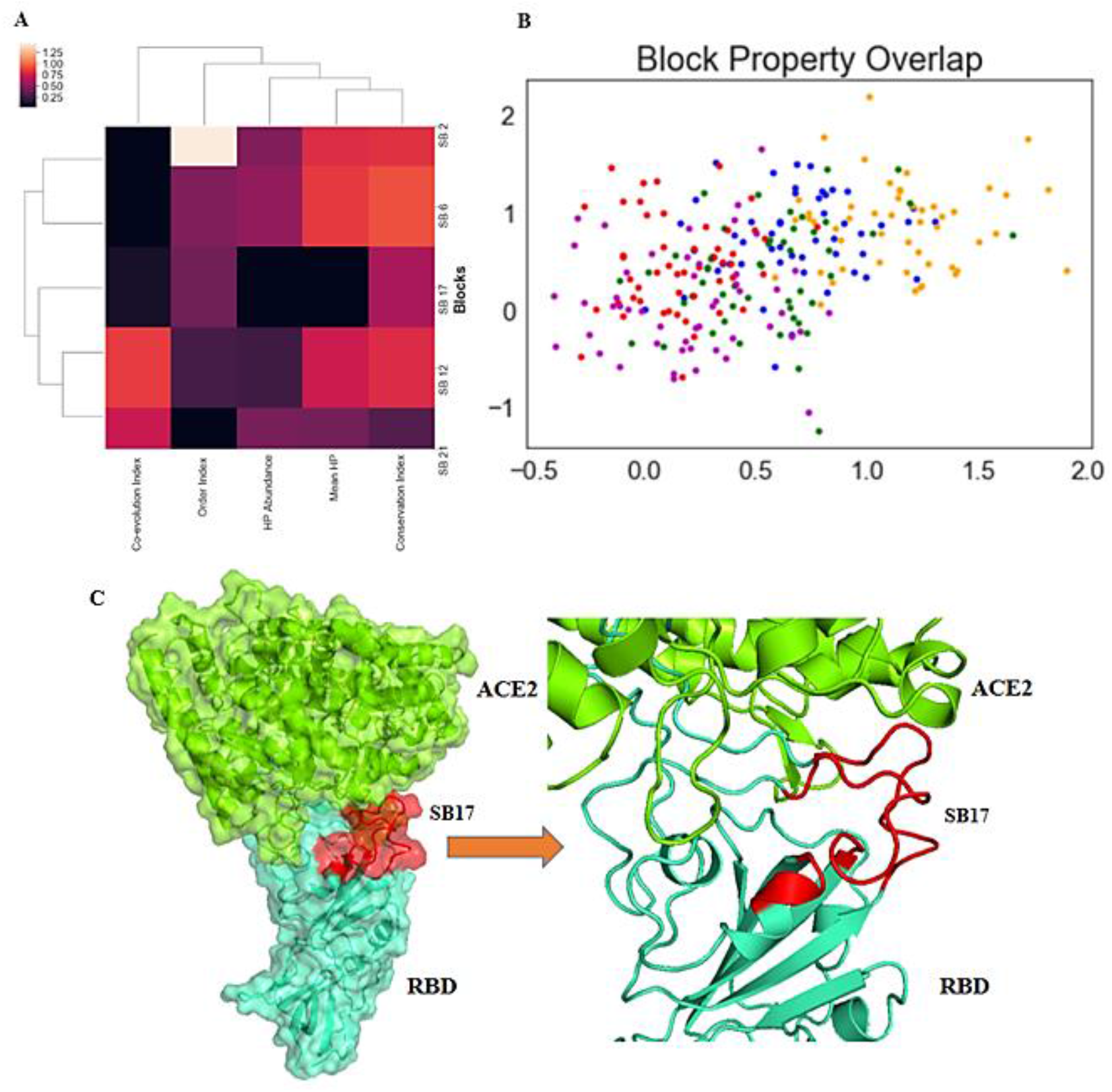
(**A**) Cluster dendrogram showing selected (from structure network analysis) structure blocks and their respective associated assigned structural and evolutionary properties. (**B**) Fuzzy clustering plot representing fuzziness in clustering of the structure blocks depending on the cumulative contribution of structural and evolutionary features. SB2 is indicated by blue sphere, SB6 is indicated by orange spheres, SB12 is indicated by green spheres, SB17 is indicated by red spheres and SB21 is indicated by magenta spheres. (**C**) In the ACE2 receptor (green region) bound RBD structure, the most important SB, SB 17 has been highlighted by red colour. Rest part of the RBD has been highlighted by cyan.

Fuzzy clustering analysis was carried out to understand, which blocks have maximal overlap and which one stands out with unique evolutionary and structural trait. We resorted to fuzzy c-means clustering technique as we believed that soft clustering approach would essentially capture overlap among the blocks and hence their evolutionary and structural traits. A considerable overlap was observed with maximum for SB2 and relatively lower for SB6 (**Figure 6B**).

## Discussion

Our current study provided a mechanistic insight into the evolutionary trail and structural facets of spike RBD resorting to an integrated evolutionary- and structure-guided approach, thereby capturing hidden uncharted traits of protein. This integrated approach cumulatively captures and explains some critical traits of the RBD with further identification of the regions key to both evolution and structural scope. Our sequence space exploration using sequence conservation and co-evolution signal provided an insight into the constraints of RBD sequence space and reflected the importance of stretch spanning from 121 to 180 (i.e., 439 to 478 in the S-protein), which strikingly coincides with the region of the protein forming the RBD-ACE2 interaction interface and which is predicted to be highly disordered. Further guided by our deep mutation analysis and epistasis model, we understood how the aforementioned stretch is evolutionarily an interplay between the conservation and variation. Deep mutation studies provided position-wide (121 to 180; i.e., 439 to 478 in the S-protein) insight into the mutational tolerance. The deep mutation landscape potentially explained variations observed in the RBD sequence space and probable variations, which could be expected during mutations in the evolutionary trajectory. This stands out extremely critical and warrants comparison with evolving virus genome, as rapid mutation fixation is a key event in the viral evolution and RBD evolution is extremely critical as RBD dictates host receptor binding and all subsequent propagation events. We referred to the closest relatives of Spike-RBD to understand the variations observed in the region 121/439 to 180/478 and their possible consequences on the spike-RBD structure. Statistical energy computed using epistasis model revealed comparable stabilities and no specific bias towards fixation of any of the substitution mutations observed. Thus, this stretch was a perfect poise of conservation and variation yet not disrupting the co-evolutionary dynamics.

Structure network analysis approach was deployed to understand the internal orchestration of RBD and how the global organization and structural adaptability is dictated by the significant SBs (which, in turn, house the majority of these evolutionarily co-evolving residues). From the community cluster network, six SBs were selected as significant by considering the number of residues housed inside (Table 1) and due to their dense connections with other clusters (**Figure 5A**). These observations also indicated their importance to the overall structural dynamics of the domain. Interestingly, majority of the residues observed to be directly involved in interaction with the receptor protein from our docking study, were found to be distributed in all the important SBs, apart from SB6 (the biggest SB in terms of number of constituent amino acids). SB6 was found to be comprised of the major number of residues with mild Shannon’s entropy values, indicating their moderate limit of tolerance to mutations (as quantified by ∆E with lower being comparable to the current wild-type form). These residues did not exhibit high propensity towards coevolution or being fully conserved. Hence, SB6, with its balanced abundance of conserved and co-evolving residues, can be hypothesized to be critical cluster, which is further validated by its constituent residues, which impose significant impact on the residue interacting with ACE2 receptor. Monte Carlo simulation revealed the importance of two sub-stretches, from residue 129/447 to 166/484 (residues of SB21) and from residue 123/441 to 127/445 (constituent of SB17). Among these two patches, the residues in SB 21 were observed to be in close vicinity with the binding groove of ACE2 receptor. Residue 123/441, 124/442, 125/443, and 127/445 (SB17) were found to be positioned in close proximity to the binding residues and in turn contributing to the conformational flexibility needed for effective binding. It is interesting to note that entire structural segment, which participates in the receptor recognition and occurs in and around the binding cleft is entirely loop-dominant and is predicted to be highly disordered. From an evolutionary and structural point of view it indicates high extent of mutational tolerance, as single substitution mutation would not impact low order structures viz. loops in this case and malleability needed to effective binding.

**Table 1:** RBD Structure Block/Cluster Members

| Structure Block/Cluster id | Residue Member |
| --- | --- |
| 1 | 319 |
| 2 | 320:325, 374:376, 399:400, 407:410, 434:436, 485, 488:489, 508:510, 540:541 |
| 3 | 326 |
| 4 | 327, 530, 532:533 |
| 5 | 328 |
| 6 | 329:330, 333:338, 355:366, 369, 377:398, 411:415, 425:433, 511:527, 534:539, 544:545, 550:555, 585:591 |
| 7 | 331:332 |
| 8 | 339 |
| 9 | 340:345 |
| 10 | 346 |
| 11 | 347:348, 354 |
| 12 | 349:353, 416, 418:424, 454, 457:469, 554:569, 571:578, 582, 584 |
| 13 | 367:368 |
| 14 | 370:373 |
| 15 | 401 |
| 16 | 402 |
| 17 | 403:406, 438:451, 496:507 |
| 18 | 317 |
| 19 | 437 |
| 20 | 452:453, 455, 492:495 |
| 21 | 456, 470:484, 486:487, 490:491 |
| 22 | 528:529 |
| 23 | 531 |
| 24 | 542 |
| 25 | 543 |
| 26 | 546 |
| 27 | 547 |
| 28 | 348 |
| 29 | 570 |
| 30 | 579:581, 583 |

Comparative analysis explored how the local structural orchestration and the stability factors, such as hydrophobicity (HP), have been associated with the evolutionary context of RBD. This comparison predicted that a SB with high HP index being comprised of majority of the hydrophobic residues demonstrated the propensity to become less co-evolving and vice versa. Hence, majority of the interacting residues of RBD were positioned in the SBs with less HP. This comparison of structural property with conservation/coevolution trend for the selected sub-blocks of the protein uncovered the significance of structural adaptability in sustaining the functional dynamics of the enzyme, along with the sequence variations that confer specificity. Our extension of the study to deploy fuzzy clustering technique was needed to have an approach, which could effectively unveil the overlap of RBD substructures (which we referred to as the structure blocks) based on their cumulative evolutionary and structural traits. We observed that the SB6 stands out with a unique distribution pattern having the least overlap owing to its distinct structural features and propensity towards conservation.

On the other hand a significant overlap in the fuzzy cluster profile was observed between SB17 and SB21 based on their evolutionary and structural features. Some residue stretches in SB17 and all residues of the SB21 were observed to be highly co-varying from evolutionary perspective as quantified from MI. Being associated with ACE2 interaction and forming the RBD-ACE2 interaction sphere, residues constituting SB17 and SB21 were thus found to be critical for interaction and the associated conformational changes. Furthermore, the mutation landscape unveiled some residues with high tolerance to be housed in SB 17 (123/441, 125/443, 126/444, and 12/445) indicating their implication on structure and evolution.

Therefore, resorting to this two (evolutionary and structural) inter-linked component-multi-tier analysis protocol, we strongly hypothesize that SB17 (Figure 6C) is an extremely critical component of the spike RBD structure, which could be an important therapeutic target. Unlike the other reports specifically pointing the residues associated with ACE2 receptor recognition, we provided an extended picture of regions in the protein structure, which, if targeted, could potentially inhibit virus propagation and block the probable mutational escape routes, thereby functioning as an anti-evolution strategy. Furthermore, this evolution- and structure-based analytical pipeline could potentially be used to understand the critical sub-structures of other proteins and could be plugged in for other therapeutic purposes.

## Materials and Methods

We started with the sequence retrieved from the recently solved Cryo-EM structure of SARS-CoV-2 spike protein (10). Here we selected only the receptor binding domain (RBD) of the protein for our study. NCBI Protein Blast was performed with the sequence corresponds to the RBD. Only those non-redundant protein sequences were selected that shares at least 75% sequence identity with the RBD region. This enzyme dataset was further used in Clustal Omega in order to obtain multiple sequence alignment (MSA) (11, 12). Followed by MSA, sequence space analysis was performed. Then protein structure was investigated in detail and connection between sequence information and structure were studied.

### Sequence Space Analysis

Phylogenetic tree was constructed using neighbor-joining (NJ) (13), algorithm based on the aligned sequence as implemented in the Rate4Site program(14).

By analyzing the MSA of homologous proteins, we can explore two types of residue information: conserved amino acids at certain positions and co-varying residues throughout the course of evolution (3, 15). Amino acids that do not alter throughout the evolutionary timeframe are designated as conserved residues. Mutations in non-conserved positions result in compensatory substitution at another position to preserve or restore the structural orchestration as well as biophysical properties of the protein. Since evolutionary perturbations in the sequences are constrained by a number of requirements, using the information confined in MSAs would have been useful to predict residues which were likely to be interdependent in the three-dimensional structure. Here we deployed information theory to predict positional correlations in MSA to investigate these conserved as well as interdependent positions respectively having structural or functional significance (3, 16).

In order to refine the MSA by reducing the number of gaps, any column in the alignment for which there was a gap in the specified coordinate file segment was removed. Similarly any rows comprised of more than 20% gaps were also removed.

To reveal evolutionary conserved positions, entropy calculation was accomplished as a function of sequence conservation. By deploying the Shannon information entropy measurement, the tolerance of a particular sequence position to amino acid substitution was understood (17).

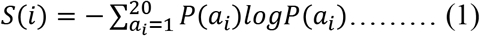

In equation 1, ‘*i*’ represented the sequence position, *P(a*_*i*_) designated the probability of amino acid ‘*a*’ to be present at the ‘*i*’th column of MSA. *S(i)* represented the Shannon entropy score, its lower values correspond to the fully conserved amino acid residues at *i*th position (16). Whereas increase in Shannon entropy score indicated the probability of that particular position to be less conserved, i.e. more random. Gaps in each column were treated as uniformly distributed amino acids (16).

Structural as well as functional restraints of a protein molecule could be labeled by the evolutionary co-varying segments (18–21). In order to understand the degree of co-evolutionary relations between amino acid positions of the RBD region, i.e., interdependency of the residues along the sequence, mutual information (MI) theory was deployed,

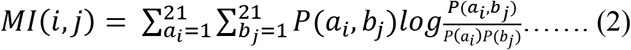

Where *P(ai, bj)* described the probability of finding amino acids of type a and b at the respective sequence positions *i* and *j* simultaneously (22). *MI(i,j)* indicated the coevolution propensity of position i and j. The gaps were treated as the 21st amino acid type. This *MI(i,j)* value diverges in the range from 0 to MI_max_ where 0 corresponds to the fully uncorrelated residues and the highest value indicate the most interdependent pairs of residues. In order to study the correlation between highly coevolving residues in RBD in detail, cut off value for MI was selected as 0.75. We went on to calculate the conservation score for individual sites which corresponds to the position’s evolutionary rate. The rate of evolution at each site was calculated using the empirical Bayesian (Mayrose et al., 2004). The stochastic process underlying the sequence evolution and the phylogenetic tree were explicitly considered.

We resorted to an unsupervised method for predicting the mutation effects that specifically captures residue dependencies between positions. On Python 3 we deployed EVcoupling to assess the quantitative effects of mutations in RBD(23). Deep mutational scan was performed to understand the effects of substitution mutations in RBD and generate the mutational landscape. Existent proteins show signatures of selection throughout their evolutionary trajectory. For genes separated by hundreds of millions of years it is not uncommon that they exhibit negligible sequence identity and still exhibit remarkable conservation of their structures and functions. We aimed to present a statistical approach that could potentially reveal dominant constraints.

In our analysis the evolutionary a sequence *σ* with probability *P(σ)* was denoted as:

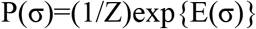

“Energy” function *E(σ)* in the model was to capture constraints on the sequences. *Z* is to normalize the distribution to sum to one over all possible sequences of a fixed length. *E(σ)* was represented as (negative) energy of a model from statistical physics or as proportional to the scaled fitness *N*_*e*_*F* in toy, equilibrium models of population genetics(23). We used the energy function *E(σ)* with two types of constraints viz. 1.pairwise constraints to depict co-dependencies in combinations of amino acids for each pair of sites and 2.site-specific constraints reflecting bias towards or away from specific amino acids at each position. As per the calculation on EVcoupling platform the total energy for a specific sequence *E(σ)* was represented sum of coupling terms **J**_*ij*_ between every pair of residues and a sum of site-wise bias terms **h**_*i*_ (fields),

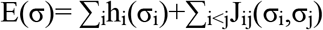

Combining the sequence model with our energy function a model was generated which is basically a pairwise undirected graphical model in computer science and a Potts model in statistical physics(23).

We used these models to make sequence-specific predictions representing the relative selective chances of mutation events. Starting from a multiple sequence alignment, we measured the site and coupling parameters **h** and **J** using regularized maximum pseudo-likelihood. After the parameters were inferred, we measured the effects of single or higher-order substitutions on a particular sequence background with the log-odds ratio of sequence probabilities between the WT form and substitution-mutant sequences.

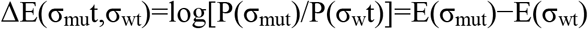

Sequence logo was generated using Seq2logo platform using aligned sequences as inputs. Logo was generated using Kullback-Leibler method to give a clear visual image of the conserved and variable regions. The Y-axis describes the amount of information in bits. The X-axis shows the position in the alignment. At each position the stack of symbols represents the amino acids observed to make up the sequence space. Large symbols represent frequently observed amino acids, big stacks represent conserved positions and small stacks represent variable positions.

Hobohm clustering algorithm was selected during logo generation which was devised to reduce redundancy of biological sequence data sets and the threshold was set at 0.63.

Co-evolutionary network was generated using Thomas Fruchterman & Edward Reingold graph layout (24). It simulates the graph as a system of mass particles in a force directed layout. The nodes are the mass particles and the edges are springs between the particles. The algorithms try to minimize the energy of this physical system.

### Structure-Based Analysis

#### Model Selection

Sequence retrieved from the recently solved Cryo-EM structure of S protein were subjected to model building using I-TASSER (25). The reported stretch corresponds to RBD, i.e. 319 to 591 were truncated from the built structure of the protein. The ACE2 receptor protein model was resorted from PDB (ID: 1R42). These models were used for further analysis.

#### Structure Network Analysis

The structure network illustration of a protein is a depiction of topological analysis of 3D structure irrespective of its secondary structure and folding type (26). The internal motions as well as structural dynamics of proteins are very much associated with its function and activity; hence we used normal mode analysis (NMA) for the prediction of functional motions in the protein segment (27). Followed by NMA, a correlation analysis was performed to generate cross-correlation matrix. Then by means of correlation network analysis, we generated structure network using the RBD of the spike protein.

By means of correlation network analysis, an all residue network was generated and it was split into a highly correlated coarse grained community cluster network by using Girvan-Newman clustering method where the highly interacting residues were clumped together in the clusters (28).

#### Molecular simulation

To understand the dynamics of the RBD we resorted to Monte Carlo simulation technique to simulate the RBD dynamics deploying CABS (C-alpha, beta, and side chain) coarse grained protein model. We deployed the standalone version CABS-flex on python 3 which employs the Monte Carlo dynamics and asymmetric Metropolis scheme satisfying the requirements of microscopic reversibility and Boltzmann distribution of generated ensembles(29). The simulation parameters were modified at the number of cycle (Ncycle) and number of model skipped keeping the seed for random number generator at 3864. The ‘Number of cycles’(Ncycle) field was set at 100 resulting in 20 × 100 = 2000 models in the trajectory The ‘Cycles between trajectory frames’ (Nskipped) which refers to the number of models skipped on saving models was kept at 100. Total numbers of generated models were thus 20×100×100= 200,000. We used a T=1.2 which is close to the native state temperature.

We further deployed Tanford-Kirkwood model (TK) in which protein molecule is treated by a spherical cavity with dielectric constant ϵ_p_ and radius b surrounded by an electrolyte solution modeled by the Debye-Hückel theory(8). We further resorted to the modification of the model which included solvent static accessibility rectification for each of the residues which are ionisable and which takes into account the irregular protein-solvent interface. The model is referred as the Tanford-Kirkwood model with solvent accessibility and we in our study would refer to it as TKSA.

#### Molecular Docking Study

In order to determine the structure of the ACE2-RBD complex and to identify the residue/segments of the RBD interacting with the ACE2 receptor protein, we resorted the computational strategy of protein-protein docking (30). In this approach, the coordinate information of the ACE2 receptor protein was resorted from protein data bank (PDB) (PDB ID: 1R42). The energy minimized model of ACE2 was subjected to interact with the model RBD structure using the tools from ClusPro depending on the PIPER docking algorithm based on the Fast Fourier Transform Correlation technique (31, 32). In this rigid body docking method, the ACE2 receptor protein was considered as the rigid body, and the RBD segment (considered as the ligand) was placed on a movable grid, where the angular step size for rotational sampling of ligand orientations was set to about 5° in terms of Eular angles. Energy minimization followed by root-mean-square deviation (RMSD) based clustering was performed for accurate and near-native conformational sampling and refinement of the complex structure (32). The topmost docking outputs defined by centers comprised of highly populated clusters correspond to lowest interaction energy between the two proteins.

#### Comparison between Sequence and Structure Index

In order to bridge evolutionary index with the structural properties of different selected significant segments of the receptor binding domain, we imposed three subsets of properties that decipher structural characteristics and two subsets decoding the sequence space information of the SBs. Here only the sub-blocks (SBs) having significant number of residues as well as significant impact on structure were focused. Three imposed structural traits were Mean Hydrophobicity (Mean HP), Hydrophobicity Abundance (HP Abundance) and Order Index. Similarly conservation and coevolution index were also calculated for the highly interdependent positional patches in the sequence.

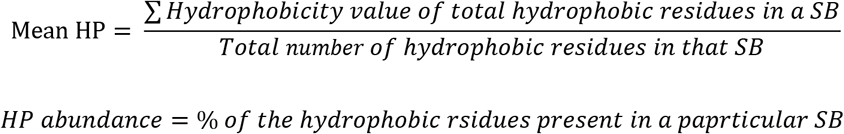

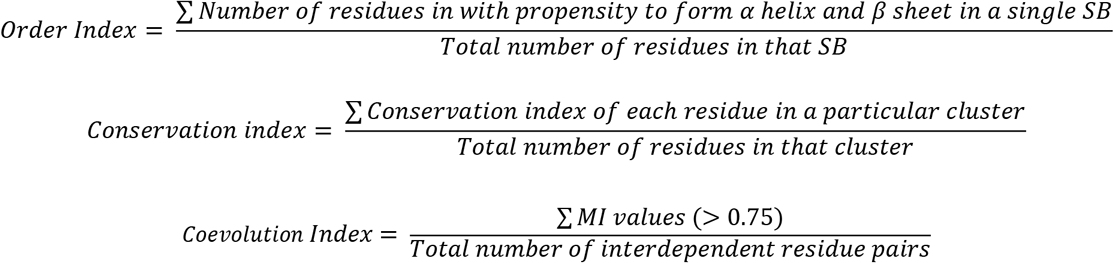

In order to understand the correlation between the structural and sequence properties within the selected significant SBs (structure blocks), a cluster dendogram was generated.

### Fuzzy C-means Clustering

Fuzzy clustering based on fuzzy logic principles was used to cluster multi-dimensional structure blocks of Spike-RBD where each block had evolutionary and structural features. FCM or fuzzy C-means clustering technique was utilized to partition a finite collection of “n” elements X={x_1_,… x_n_} into a collection of c fuzzy clusters. With a finite set of data, the algorithm returns a list of c cluster centres as C={c_1_……c_n_) along with a partition matrix. Any point x has a set of coefficients giving the degree of being in the kth cluster wk(x). With fuzzy c-means, the centroid of a cluster is the mean of all points, weighted by their degree of belonging to the cluster, or, mathematically,

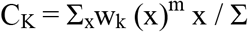

Where, m is the hyper-parameter that controls how fuzzy the cluster will be. The higher it is, the fuzzier the cluster will be in the end.

### Analysis and Representation

Majority of evolutionary and structural analysis were done with Python3. Visual renditions were made using Seaborn library of Python. For statistical analysis and representations Origin Pro 9.0 and Tableau were also used. For Graph theoretical modelling Gephi graphing tool was used. Protein models were represented using PyMol. Structure network analysis using Bio3D package was carried out on RStudio.

## Acknowledgment

DS acknowledges Department of Science and Technology (DST), Govt. of India for doctoral fellowship (DST-INSPIRE). DS and KC acknowledge the Director, IICB.

## Author contribution statement

SC conceived the project. KC and SC planned the overall project outline. DS and SC performed the computational experiments and analysis. KC and VU critically analyzed the manuscript and VU performed IDP analysis.

## Declarations of interest

Authors don’t have any conflict of interest.

## Notes

### Competing Interest Statement

The authors have declared no competing interest.

## References

1. Jaimes JA, André NM, Chappie JS, Millet JK, & Whittaker, GRJJoMB (2020) Phylogenetic Analysis and Structural Modeling of SARS-CoV-2 Spike Protein Reveals an Evolutionary Distinct and Proteolytically-Sensitive Activation Loop.

2. Uversky, VN (2015) Unreported intrinsic disorder in proteins: disorder emergency room. Intrinsically disordered proteins 3(1):e1010999.

3. Chowdhury S, et al. (2019) Evolutionary Analyses of Sequence and Structure Space Unravel the Structural Facets of SOD1. 9(12):826.

4. Chowdhury S, Banerjee A, & Chattopadhyay K (2017) Metal ion co-factors sculpt the heterogeneity of conformational landscape in Superoxide Dismutase. EUROPEAN BIOPHYSICS JOURNAL WITH BIOPHYSICS LETTERS, (SPRINGER 233 SPRING ST, NEW YORK, NY 10013 USA), pp S344–S344.

5. Sarkar-Banerjee S, et al.(2016) The non-native helical intermediate state may accumulate at low pH in the folding and aggregation landscape of the intestinal fatty acid binding protein. Biochemistry 55(32):4457–4468.

6. Alexiev U, Rimke I, & Pöhlmann T (2003) Elucidation of the nature of the conformational changes of the EF-interhelical loop in bacteriorhodopsin and of the helix VIII on the cytoplasmic surface of bovine rhodopsin: a time-resolved fluorescence depolarization study. Journal of molecular biology 328(3):705–719.

7. Ortega JT, Serrano ML, Pujol FH, & Rangel HR (2020) Role of changes in SARS-CoV-2 spike protein in the interaction with the human ACE2 receptor: An in silico analysis. EXCLI journal 19:410.

8. Contessoto VG, de Oliveira VM, Fernandes BR, Slade GG, & Leite VB (2018) TKSA-MC: A web server for rational mutation through the optimization of protein charge interactions. Proteins: Structure, Function, and Bioinformatics 86(11):1184–1188.

9. Ibarra-Molero B, Loladze VV, Makhatadze GI, & Sanchez-Ruiz JM (1999) Thermal versus guanidine-induced unfolding of ubiquitin. An analysis in terms of the contributions from charge-charge interactions to protein stability. Biochemistry 38(25):8138–8149.

10. Wrapp D, et al.(2020) Cryo-EM structure of the 2019-nCoV spike in the prefusion conformation. 367(6483):1260–1263.

11. Sievers F, et al.(2011) Fast, scalable generation of high-quality protein multiple sequence alignments using Clustal Omega. 7(1).

12. Sievers F & Higgins DGJPS (2018) Clustal Omega for making accurate alignments of many protein sequences. 27(1):135–145.

13. Saitou N & Nei M (1987) The neighbor-joining method: a new method for reconstructing phylogenetic trees. Molecular biology and evolution 4(4):406–425.

14. Pupko T, Bell RE, Mayrose I, Glaser F, & Ben-Tal N (2002) Rate4Site: an algorithmic tool for the identification of functional regions in proteins by surface mapping of evolutionary determinants within their homologues. Bioinformatics 18(suppl_1):S71–S77.

15. Chowdhury S, et al.(2019) Evolutionary Analyses of Sequence and Structure Space Unravel the Structural Facets of SOD1. Biomolecules 9(12):826.

16. Liu Y, Bahar IJMb, & evolution (2012) Sequence evolution correlates with structural dynamics. 29(9):2253–2263.

17. Cover TM & Thomas JA (2012) Elements of information theory (John Wiley & Sons).

18. Göbel U, Sander C, Schneider R, Valencia AJPS, Function,, & Bioinformatics (1994) Correlated mutations and residue contacts in proteins. 18(4):309–317.

19. Shindyalov I, Kolchanov N, Sander CJPE, Design, & Selection (1994) Can three-dimensional contacts in protein structures be predicted by analysis of correlated mutations? 7(3):349–358.

20. Atchley WR, et al. (2000) Correlations among amino acid sites in bHLH protein domains: an information theoretic analysis. 17(1):164–178.

21. Lockless SW & Ranganathan RJS (1999) Evolutionarily conserved pathways of energetic connectivity in protein families. 286(5438):295–299.

22. Dunn SD, Wahl LM, & Gloor GBJB (2008) Mutual information without the influence of phylogeny or entropy dramatically improves residue contact prediction. 24(3):333–340.

23. Hopf TA, et al.(2019) The EVcouplings Python framework for coevolutionary sequence analysis. Bioinformatics 35(9):1582–1584.

24. Fruchterman TM & Reingold EM (1991) Graph drawing by force-directed placement. Software: Practice and experience 21(11):1129–1164.

25. Roy A, Kucukural A, & Zhang YJNp (2010) I-TASSER: a unified platform for automated protein structure and function prediction. 5(4):725.

26. Linding R, et al.(2003) Protein disorder prediction: implications for structural proteomics. 11(11):1453–1459.

27. Alexander MD, et al.(2002) “True” sporadic ALS associated with a novel SOD-1 mutation. 52(5):680–683.

28. Srinivasan E & Rajasekaran RJTpj (2019) Computational investigation on electrostatic loop mutants instigating destabilization and aggregation on human SOD1 protein causing amyotrophic lateral sclerosis. 38(1):37–49.

29. Kurcinski M, et al.(2019) CABS-flex standalone: a simulation environment for fast modeling of protein flexibility. Bioinformatics 35(4):694–695.

30. Sarkar-Banerjee S, et al.(2018) The Role of Intestinal Fatty Acid Binding Proteins in Protecting Cells from Fatty Acid Induced Impairment of Mitochondrial Dynamics and Apoptosis. Cellular Physiology and Biochemistry 51(4):1658–1678.

31. Kozakov D, et al.(2013) How good is automated protein docking? 81(12):2159–2166.

32. Kozakov D, et al.(2017) The ClusPro web server for protein–protein docking. 12(2):255.

